# Ephaptic coupling between olfactory receptor neurons is sensitive to relative stimulus timing: Implications for odour source discrimination

**DOI:** 10.1101/2023.11.27.568881

**Authors:** Georg Raiser, C. Giovanni Galizia, Paul Szyszka

## Abstract

Insect olfactory receptor neurons (ORNs) are often co-localized within sensilla and exhibit non-synaptic reciprocal inhibition through ephaptic coupling. It has been postulated that this inhibition aids odour source discrimination, as synchronous arrival of different odour molecules (odorants) from a single source should increase ephaptic inhibition, whereas asynchronous arrival of odorants from different sources should decrease ephaptic inhibition. However, it was as yet unknown whether temporal arrival patterns of different odorants indeed modulate ephaptic inhibition, since past studies have focused on ephaptic inhibition of sustained ORN responses to prolonged and constant odour stimuli. However, most natural odour stimuli are not constant but rather transient and fluctuate as a result of dispersion in turbulent plumes in the air. To investigate the role of ephaptic inhibition in olfaction within turbulent environments, we recorded co-localized ORNs in the fruit fly *Drosophila melanogaster* exposed to dynamic odorant mixtures. We found that ephaptic inhibition does modulate transient ORN responses, and the strength of ephaptic inhibition decreases as the synchrony between arriving odorants decreases. These results support the hypothesis that ephaptic inhibition aids odour source discrimination.

## Background

Ephaptic coupling is a form of communication between neurons via direct electrical interaction that does not involve chemical or electrical synapses [1]. Ephaptic coupling requires close electrical proximity among neurons, achieved by sharing an electrically insulated extracellular space [1]. Such proximity is typical for insect olfactory receptor neurons (ORNs): ORNs are housed in sensilla (Fig. 1c), where multiple ORNs share the same extracellular space and are electrically insulated from the rest of the body [2].

**Figure 1.**
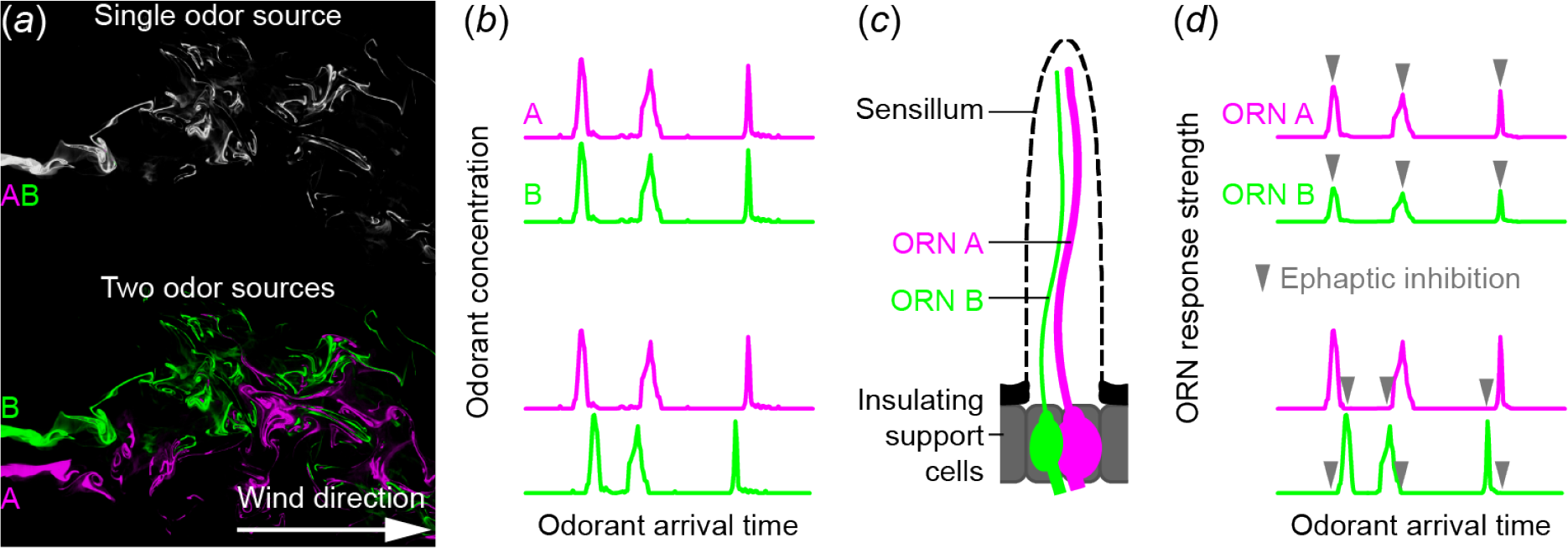
Hypothesis: Ephaptic inhibition is sensitive to relative stimulus timing and aids odour source discrimination. *Top*: Different odorants (A and B) from a single source (*a*) arrive synchronously (*b*) at olfactory receptor neurons (ORN) (*c*). Co-localized ORNs inhibit each other’s response via ephaptic inhibition (*d*), making the odorant mixture’s neural representation different from the sum of its parts A and B (synthetic odour processing). *Bottom*: Odorants (A and B) from different sources (*a*) arrive asynchronously (*b*) at the ORNs (*c*). Consequently, ephaptic inhibition is active when the affected ORN exhibits weak or no response (*d*). This results in the neural representation of the odorant mixture resembling the sum of its individual components, A and B (analytical odor processing). Data in (*a*) from [9].

Electrical modelling has predicted that co-localized insect ORNs mutually inhibit each other through ephaptic coupling [3], and such inhibition was experimentally demonstrated in fruit flies [4–6]: When an ORN *A* is continuously stimulated with its cognate odorant A (which does not activate a co-localized ORN *B*), and ORN *B* is stimulated with a pulse of its cognate odorant B (which does not activate ORN *A*), then the sustained response of ORN *A* is inhibited by the response of ORN *B*.

It has been hypothesized that fast and temporally precise ephaptic inhibition between ORNs enables insects to detect whether different odorants originate from the same or from different sources (Fig. 1) [7]. This hypothesis was motivated by the following observation in corn earworm moths (*Helicoverpa zea*) [7]: Male moths take off for a search flight when they encounter packets of a female moth’s sex pheromone component (A), even when A is mixed with sex pheromone packets (B) of another moth species. However, when A and B are released from the same source, the male moths do not take off. This behavioural inhibition has been explained by the synchrony between A and B, which occurs when A and B originate from the same source (Fig. 1a, b) – indicating that the source is a female moth from a different species. Ephaptic inhibition could enhance this discrimination between synchrony and asynchrony: In corn earworm moths, ORNs tuned to own and alien pheromone components are co-localized in a single sensillum (ORN A and ORN B in Fig. 1c). The mutual inhibition between ORN A and ORN B would become apparent when A and B arrive synchronously (Fig. 1d), and the inhibited response of ORN A would then fail to trigger a search flight. A recent study modelled such an odour source discrimination mechanism [8].

For this source discrimination mechanism to work, we predict that in a turbulent stimulus situation, ephaptic inhibition would affect the transient ORNs’ responses to the onset of an odour stimulus, and the strength of synaptic inhibition would decrease with increasing asynchrony between arriving odorants. However, this has, to our knowledge, not yet been shown experimentally.

Here, we tested whether ephaptic inhibition affects transient ORN responses to odorant onset and to fluctuating odorant stimuli by recording a pair of ORNs within one fruit fly sensillum (the ab3A and ab3B ORNs). We chose the ab3 sensillum since it is known that ab3A and ab3B inhibit each other’s sustained responses to constant odorant stimuli via ephaptic coupling [6].

Our results show that ephaptic inhibition strength is modulated by the relative timing of odorant onsets, and thus could support odour source segregation in a natural environment: Mixtures of synchronously or correlated fluctuating odorant stimuli elicit ephaptic inhibition, whereas mixtures of asynchronously or uncorrelated fluctuating odorant stimuli elicit weak or no ephaptic inhibition. The synchrony window of ORN’s ephaptic inhibition is approximately 50 ms wide (range of onset asynchrony that induces ephaptic inhibition), aligning with the ∼ 30 ms synchrony window observed in behavioural experiments [10].

## Methods

### Single sensillum recordings in *Drosophila*

To measure the activity of multiple neurons within a single sensillum, we performed single sensillum recordings [11] and recorded from large basiconic ab3 sensilla of female *Drosophila melanogaster* wild type Canton S flies, aged 1 - 9 days. The flies were raised at 25 °C on a standard *Drosophila* medium, with a 12/12 h day/night cycle. Recordings were performed during the flies’ light period. Flies were immobilized in a pipette tip and mounted ventral side facing upwards on a custom 3D printed holder. The antennae were bent outwards from the head and stabilized by a small droplet of n-eicosane (Sigma Aldrich) onto the edge of a microscopy glass slide. Sensilla were localized using a 50x magnification lens (50x/0.55 DIC EC Epiplan-Neofluar, Zeiss) with brightfield illumination. The recording and reference electrodes were tungsten wires (diameter = 0.1mm), which were electrolytically sharpened with AC-current in a 0.5 M KOH solution. The recording electrode was inserted into the sensillum with a micromanipulator (Sensapex, SMXS-K-R). The reference electrode was inserted into the eye. The voltage difference between sensillar lymph and hemolymph (the transepithelial potential) was amplified x1000 with an MA 103 headstage and MA 102 amplifier (Universitaet zu Koeln) in DC mode, bandpass-filtered between 1 and 8,000 Hz. Noise from the powerline was reduced by a Hum Bug (Quest Scientific). Signals were digitized using a Micro 3 1401 A/D converter (Cambridge Electronic Design). Sensillar identity was determined by applying a sequence of diagnostic odorants that unambiguously identify the sensillar identity (ethyl hexanoate, methyl acetate, isobutyl acetate, 2-butanone, 2,3-butanedione, 1-hexanol, derived from the DoOR database [12]).

### Odorant stimuli

We used the odorants methyl hexanoate (MHXE, CAS: 106-70-7) and 2-heptanone (HEPN, CAS: 110-43-0) (Sigma Aldrich). Odorant stimuli were generated with a custom-made stimulator optimized for temporal precision in binary mixtures [13]. We upgraded the published version of this stimulator by an air dilution system (Fig. S1a) which eliminated a decrease in odorant concentration within and across repeated stimulations (Fig. S1b). Odorants were either diluted by a factor 10^−2^ for “high concentration” or by a factor 10^−5^ for “low concentration”. In this study, odorant stimuli were either brief pulses of odorants added into a longer-lasting background odorant, pulses with varying onset delays in the millisecond range, or randomly fluctuating odorant concentrations. Reproducible streams of fluctuating odorant concentrations (Fig. 4) were generated by switching the stimulator’s valves at random time points, where the switching times were generated from a Poisson point process with mean interval of 50 ms for a 10 s long segment. Two sequences of valve state changes for both odorants A and B were calculated separately. From those sequences, different correlated and uncorrelated mixture stimuli were constructed: For the uncorrelated mixtures, each of the stimuli A and B followed its own independent random sequences (abbreviated AB_i_). For the correlated mixtures both odorants A and B followed the identical time course, either of A (AB_A_) or of B (AB_B_).

### Data analysis

Spike sorting was performed with a custom spike sorting algorithm for single sensillum recordings [14]. All further data analysis was performed with custom Python scripts.

Spike rate estimation was performed by convolving the estimated spike trains with an alpha kernel with a standard width of σ = 20 ms. An alpha kernel ensured that the predicted spike rates do not rise before an actual spike occurs [15].

Spike train dissimilarity was quantified as Victor-Purpura distance [16]. Briefly, the VP distance between two spike trains is calculated by transforming one spike train into the other, and each transformation (removal of a spike, addition of a spike or shift by a defined time) is associated with a cost. The algorithm then finds the lowest cost to transform one spike train into the other. VP was computed with the function *spike_train_dissimilarity*.*victor_purpura_dist()* of the python module *elephant*.

## Results

First, we confirmed ephaptic inhibition between co-localized ORNs in ab3 sensilla, which contain two ORNs, ab3A and ab3B [11]. ab3A is larger than ab3B and can be recognized in single sensillum recordings by its larger spikes (Fig. 2a). We used the two odorants, methyl hexanoate (MHXE) and 2-heptanone (HEPN) [6]. MHXE activates ab3A (cognate odorant for ab3A) and not ab3B (incognate odorant for ab3B), while HEPN activates ab3B (cognate odorant for ab3B) and not ab3A (incognate odorant for ab3A).

**Figure 2.**
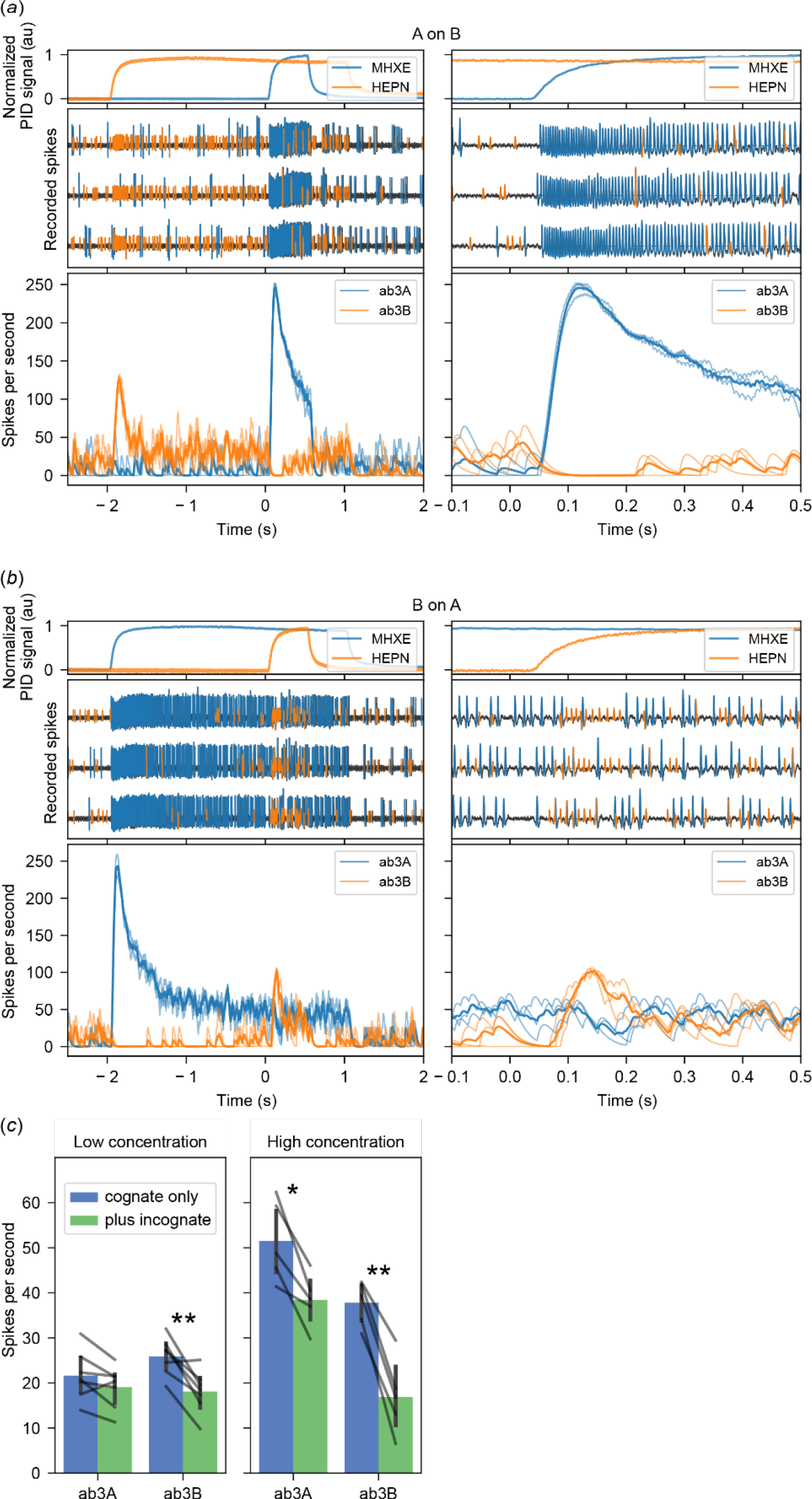
Confirmation of ephaptic inhibition of sustained ORN responses to constant odorant stimuli. (*a*) *Top*: Time course of the two odorant stimuli, MHXE and HEPN, measured with a photoionisation detector (PID). MHXE is the cognate odorant for ab3A, and HEPN is the cognate odorant for ab3B. A pulse of MHXE was introduced into a constant background of HEPN, both at high concentrations. The moment the valve opened for the MHXE pulse is marked as Time = 0. The right panels provide a detailed view of the left. *Middle*: Three recordings from individual ab3 sensilla in 3 flies during stimulation with high-concentration odorants. Spikes were automatically detected, with spikes from ab3A marked in blue and those from ab3B marked in orange. *Bottom*: Instantaneous spike rates of three ab3A (thin blue lines) and ab3B (thin orange lines) along with their means (bold lines). (*b*) Same as *a*) but the two odorants were interchanged. (*c*) Spike rates, recorded during 0.1 to 0.35 s after opening of the valve for the cognate odorant, are shown without (blue) and with the incognate odorant (green). Seven flies for low odorant concentrations and five flies for high concentrations. Bars show mean, and error bars show 95% confidence intervals. Stars show significance level of paired t-tests (*, p<0.05; **, p<0.005).

Adding MHXE into a HEPN background led to a decreased HEPN response in ab3B (Fig. 2a). Conversely, adding HEPN into an MHXE background led to a decreased MHXE response in ab3A (Fig. 2b), confirming reciprocal ephaptic inhibition between the two neurons [6]. Next, we investigated the effect of odorant concentration (Fig. 2c) and confirmed that ephaptic coupling increases with odorant concentration (and, accordingly, response strength) [6].

To test whether ab3A and ab3B inhibit each other’s transient response to odorant onset and whether relative onset timing between two odorants modulates ephaptic inhibition strength, we presented binary mixtures of MHXE and HEPN with shifted onsets between the two odorants (0, 3, 6, 12, 24, 48 or 96 ms onset asynchrony, with either of the two odorants as the trailing one) (Fig. 3a, b). This range covers the behaviourally relevant range, as behavioural experiments have demonstrated that fruit flies can detect odorant onset asynchronies as brief as 33 ms [10].

**Figure 3.**
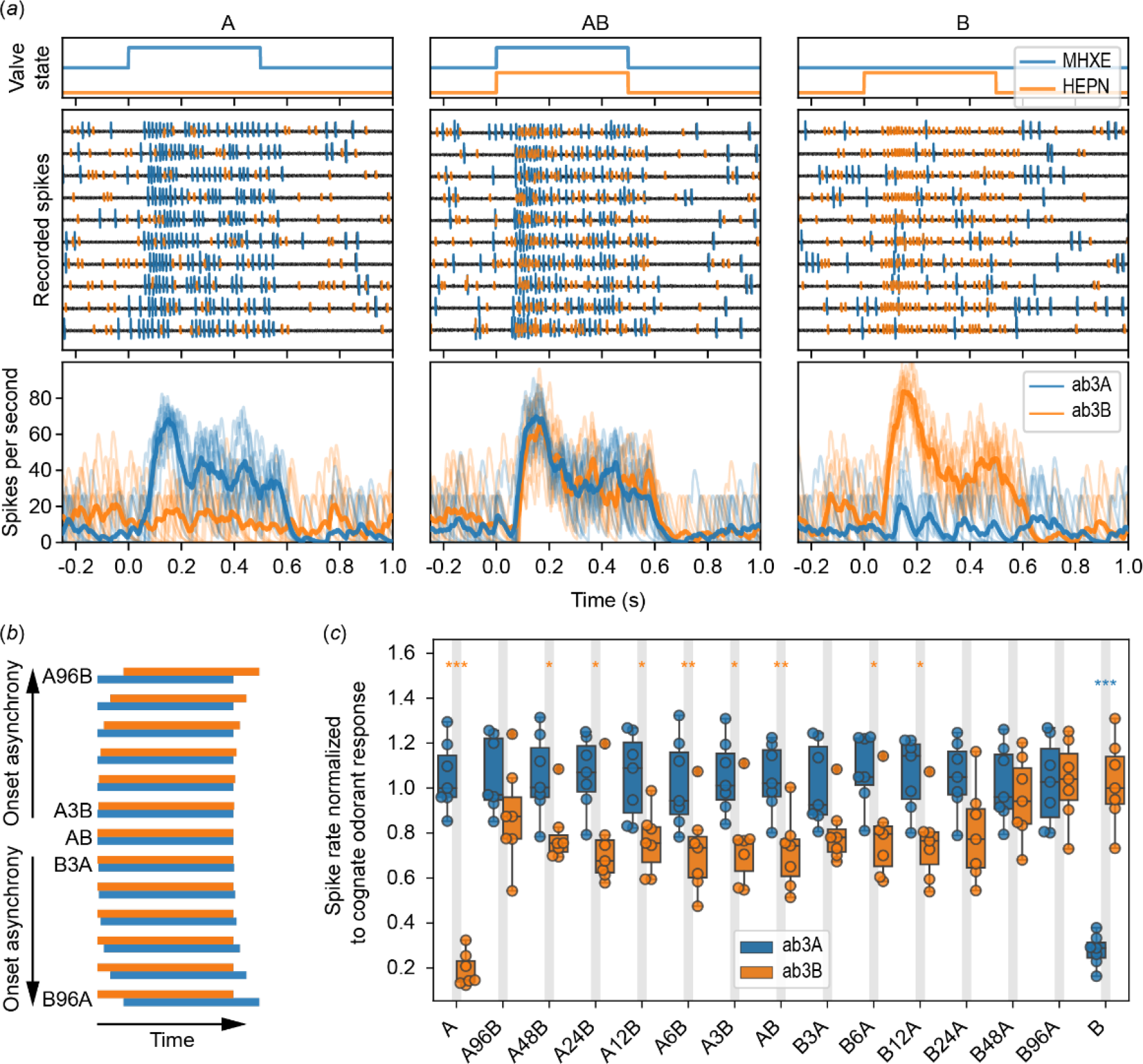
Relative onset timing in the millisecond range modulates ephaptic inhibition strength. (*a*) *Top*: Valve states for delivering 0.5-seconds long pulses of MHXE (blue) and HEPN (orange). *Middle*: Recordings from an ab3 sensilla in ten trials to low-concentrated odorants. Spikes from ab3A (blue) and ab3B (orange). *Bottom*: Instantaneous spike rates of ab3A (thin blue lines) and ab3B (thin orange lines) along with their means (bold lines). The moment the valve opened for the odorant pulses is marked as Time = 0.0. (*b*) True to scale temporal patterns of 0.5-second-long pulses of MHXE (blue, odorant A) and HEPN (orange, odorant B) in mixtures with varying stimulus onset asynchronies. (*c*) Transient responses, quantified as response peak rate, of ab3A and ab3B for all stimuli at low concentration across seven flies. Note the decrease in inhibition, indicated by a higher firing rate, as onset time differences increase. Circles show individual data points, horizontal lines show medians, boxes show interquartile ranges, and whiskers extend to ±1.5 times the interquartile ranges. Stars show significance levels for comparison between ab3A and ab3B response strengths for each stimulus (two tailed t-test, corrected for false discovery rate after [17]. *, p<0.05; **; p<0.05; ***p<0.005).

ORNs ab3A and ab3B gave reliable responses to these mixtures. However, the response of ab3B to the binary mixture was reduced as compared to the response to its ligand HEPN alone. The magnitude of this reduction varied with odorant-onset mismatch: with A-to-B onset asynchronies ranging from - 48ms (A48B) to 12ms (B12A) ephaptic inhibitions were significant. However, no inhibition was evident for longer stimulus asynchronies (A96B and B24A or greater). The timing of ORNs’ response onset was not affected by ephaptic inhibition (Fig. S2).

In the experiments conducted so far, all odorant stimuli were square pulses, meaning there was a sudden increase of the odorant concentration from zero to either low or high concentration. However, naturally occurring odorant stimuli exhibit complex temporal fluctuations in concentration. Importantly, odorants from different sources have uncorrelated temporal structures, while odorants (i.e., different chemicals in a blend) from a single source have correlated temporal structures (Fig. 1a) [9,18–21]. To investigate how stimulus correlation in a realistic environment influences ephaptic coupling, we compared ORN responses to mixtures where the streams of odorants A (MHXE) and B (HEPN) were either uncorrelated and independent (AB_i_) or completely correlated and synchronous (AB_A_ or AB_B_; the index indicates whether the shared time course of A and B corresponds to the component A or B in the AB_i_ stimulus) (Fig. 4). Hence, AB_A_ and AB_B_ are two temporally different stimuli that share the same degree of temporal correlation, and AB_i_ is a stimulus in which the individual components share no temporal correlation, and all stimuli share the same amount of overall odorant concentrations.

**Figure 4.**
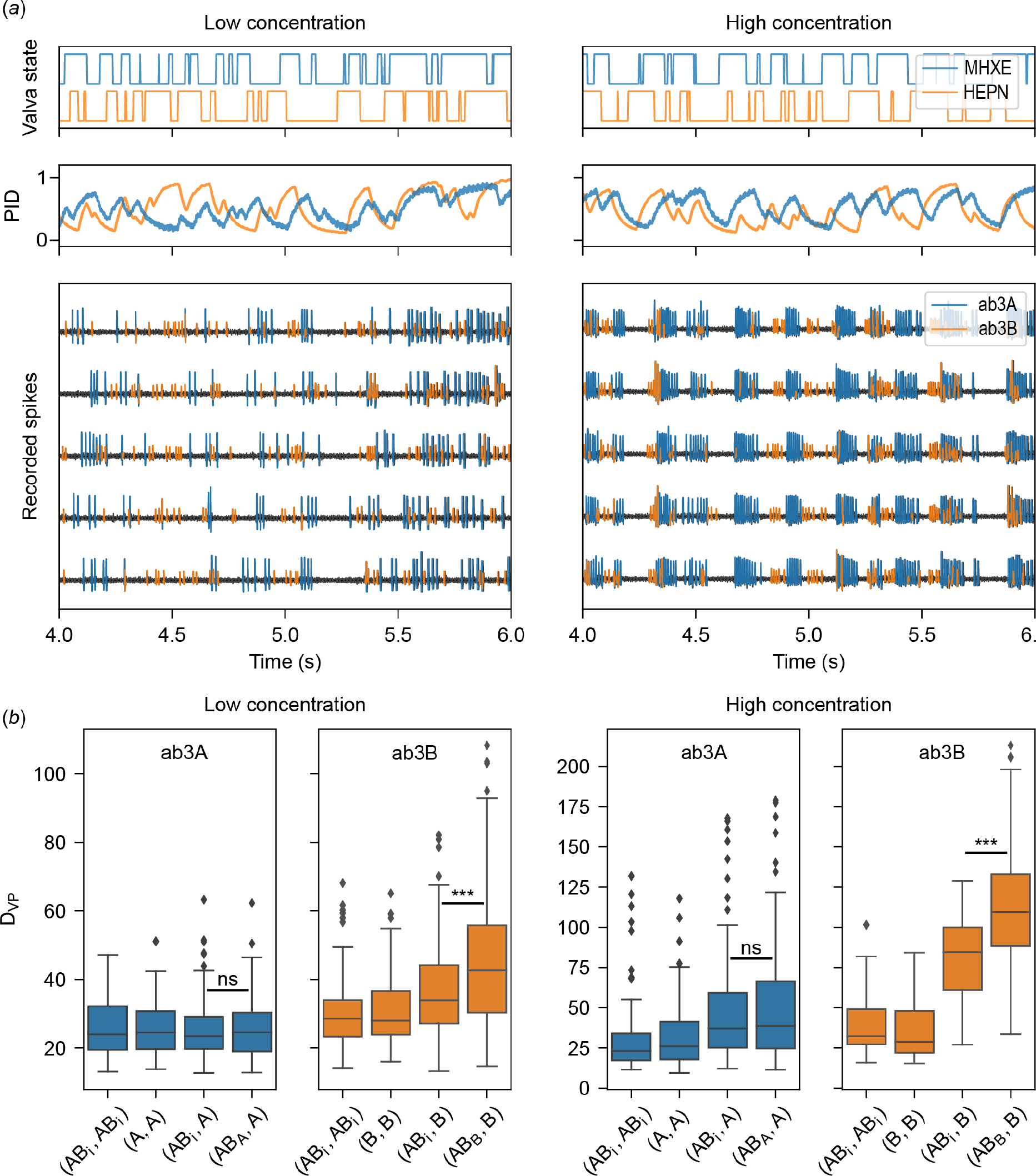
Correlation between fluctuating odorant stimuli modulates ephaptic inhibition strength. (*a*) *Top*: Valve states for delivering mixed, but uncorrelated fluctuating streams of odorant A (MHXE, blue) and B (HEPN, orange). *Middle*: Time course of the two odorant stimuli A and B measured with a photoionisation detector (PID). *Bottom*: Recordings from an ab3 sensillum in five different trials with automatically detected ab3A spikes (blue) and ab3B spikes (orange). (*b*) Victor-Purpura spike train distances (D_VP_) for different comparisons of correlated and uncorrelated odorant mixture stimulations. Box plots show the distribution of within-animal pairwise comparisons for different stimulus pairs of single odorants (A or B), correlated mixtures (AB_A_ or AB_B_; the index indicates whether A and B fluctuated with the time course A or B), and uncorrelated mixtures (AB_i_). Low values indicate similar spike trains, higher values indicate different spike trains. For example, the response to HEPN in ab3B (orange) differs between stimulus AB_B_ and stimulus B, even though the stimulus of its cognate ligand HEPN has the identical time course (right orange box, both at low and high odorant concentration). The larger difference between spike sequences between AB_B_ and B as compared to AB_i_ and B indicates that A-driven activity in ab3A exerts stronger ephaptic inhibition onto ab3B when the two odorants have synchronous time courses. Horizontal lines mark the medians, boxes show interquartile ranges, whiskers extend to ±1.5 times the interquartile ranges, and outliers are drawn as points. Stars show significance levels (two-sided U test; ***p<0.005).

Fluctuating odorant stimuli elicited reproducible spike sequences in both neurons (Fig. 4a). To measure the effect of an incognate odorant on the spike sequence, we computed Victor-Purpura distances (D_VP_) [16] between the individual spike trains. In the absence of ephaptic interactions, responses in ab3A to A, AB_i_ and AB_A_ would be equal, i.e., Victor-Purpura distance for the comparison (AB_i_, A) and (AB_A_, A) should not differ from the comparison (AB_i_, AB_i_) or (A, A). Similarly, for responses in ab3B the comparison (AB_i_, B) and (AB_B_, B) should not differ from (AB_i_, AB_i_) or (B, B). However, if an incognate odorant would create ephaptic inhibition, then the spike sequences of an ORN should differ more between its cognate odorant and the correlated mixture [(AB_A_, A) for ab3A or (AB_B_, B) for ab3B] than between its cognate odorant and the uncorrelated mixture AB_i_ [comparison (AB_i_, A) for ab3A or (AB_i_, B) for ab3B]. This is because weaker ephaptic interactions should occur if the activity peaks of both neurons do not coincide (Fig. 1d).

In both ORNs of the sensillum, there was ephaptic interaction (Fig. 4b). For example, at high odorant concentration, in ab3A, D_VP_ was larger for (AB_A_, A) than for (A, A), and in ab3B, correlated fluctuations led to more interaction than non-correlated fluctuations (Fig. 4b, D_VP_ was larger for (AB_B_, B) than for (AB_i_, B)). However, this effect was absent in ab3A. This supports the hypothesis that ephaptic inhibition is sensitive to the correlation between odour streams. The lack of this effect in ab3A could be due to its larger size in comparison to ab3B, which might result in an ephaptic inhibition from ab3B on ab3A that is too weak to be detected [4].

## Discussion

We tested Baker and colleagues’ hypothesis [7] that ephaptic inhibition between co-localized olfactory receptor neurons (ORNs) decreases with decreasing stimulus onset synchrony or correlation between fluctuating odorant streams, a mechanism that aids source discrimination and olfactory figure-ground segregation (Fig. 1) [22,23]. To this end, we first confirmed previous studies which demonstrated that in the ab3 sensillum of *Drosophila melanogaster* the response of the ORN ab3A to its cognate odorant MXHE inhibits a sustained responses of ab3B to a constant odorant stimulus of HEPN, and vice versa (Fig. 2) [4,6]. Because these studies showed that this inhibition is caused by ephaptic coupling, and we measured the same ORNs and used the same pair of odorants, we conclude that the lateral inhibition that we found is also due to ephaptic inhibition.

Our key finding is that ephaptic inhibition also affects transient ORN responses when odorant onsets are synchronous (Fig. 3) and when odorant stimuli fluctuate (Fig. 4), expanding upon previous observations of ephaptic inhibition affecting sustained ORN responses to constant odorant stimuli [4– 6]. This finding is relevant, because constant and square odorant stimuli are rarely found in natural environments, where odorants fluctuate rapidly [19,24]. Furthermore, neural responses to odorant onset flanks were found to be more informative about the identity and concentration of an odorant than sustained responses [25–32], and during odour-guided navigation, insects use the timing of odorant onsets to locate odour sources [33–42]. Therefore, our finding of ephaptic inhibition of transient ORN responses to odorant onsets suggests that this lateral inhibition plays a role in extracting information about odour identity, concentration, and odour source location. This emphasizes the significance of neuronal compartmentalisation and raises the question about the evolutionary pressures that led to the association of specific ORNs into discrete sensilla [5,8,43].

Importantly, the strength of ephaptic inhibition between ORNs varied with relative odorant onset timing (Fig. 3c), as predicted by [7]: Ephaptic inhibition was weakest with asynchronous and uncorrelated mixtures. In a natural setting, this would occur when two separate odorant sources create overlapping but desynchronized odorant plumes downwind (Fig. 1a), for example, when a female moth releases sex pheromone, and a sympatric female from another species releases her (different) pheromone from nearby [7]. The reduction of ephaptic interactions in this scenario allows for a more efficient signal separation by the male moth, helping the male to identify its conspecific female.

In addition to discriminating between the synchronous and asynchronous arrival of different pheromones, the temporal sensitivity of ephaptic inhibition could aid general odour source discrimination. Odorants from one source arrive at the antenna synchronously and as correlated odorant streams [18,19]. Those one-source odorants would induce ephaptic inhibition, leading to distinct ORN responses compared to the single odorants. This would facilitate synthetic odour processing, such that the odorants would be perceived as one unitary object (a perfume, or gestalt) and thus interpreted as coming from a single source. However, odorants from two different sources arrive at the antenna asynchronously and as uncorrelated odorant streams (Fig. 1) [9,18–21]. Those two-source odorants would induce less ephaptic inhibition, leading to similar ORN responses compared to the single odorants. This would facilitate analytical odour processing, such that the odorants would be perceived as a mixture of two distinct and recognizable compounds - or as coming from two separate sources.

Our finding that the synchrony window triggering ORNs’ ephaptic inhibition (i.e., the -48 to 12 ms onset asynchrony, Fig. 3c) comprises the 33 ms-synchrony window identified for odour source discrimination in behavioural experiments [10] strengthens the hypothesis that ephaptic inhibition aids this discrimination. This hypothesis finds further support in studies demonstrating co-localisation of ORNs responsive to odorants with opposing valences [5,43–47], and in studies demonstrating that behavioural responses to attractive odorants diminish when antagonistic odorants arrive synchronously but not when they arrive asynchronously [10,22,45,48–50]. Exploring the temporal sensitivity of ephaptic inhibition between ORNs in the same insect species as in these studies will provide further insights in its functional role in odour source discrimination.

## Authors’ contributions

Conceptualization, CGG, GR, PS; Methodology, GR; Formal Analysis, GR; Investigation, GR; Resources, CGG, PS; Data Curation, GR; Writing—Original Draft, GR, PS; Writing— Review & Editing, CGG, GR, PS; Funding Acquisition, CGG, PS. All authors read and approved the final manuscript.

## Supplementary Materials

**Figure S1.**
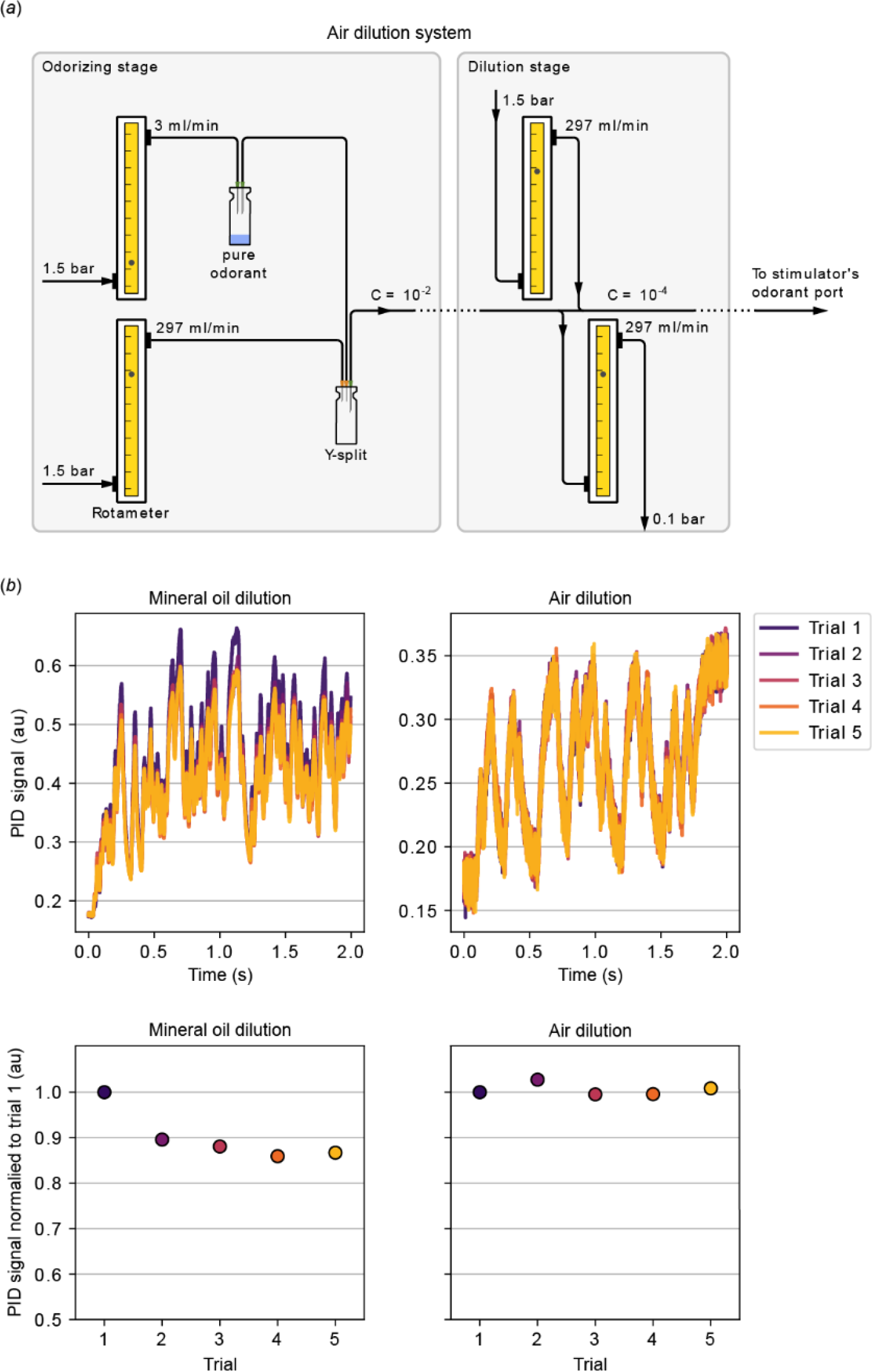
Air dilution system for stable odorant concentrations. (*a*) Pressure-controlled (35898; Analyt-MTC), charcoal-filtered air from in-house air supply was dived in a 1:100 ratio (3 and 297 ml/min) with flowmeters (112-02GL, Analyt-MTC). After the smaller fraction was routed through the headspace of a 20 ml glass vials (Schmidlin, sealed with a Teflon septum), both flows were combined. This served as an initial 10^−2^ dilution. Further dilutions were generated by first removing 297 ml/min, and subsequently adding 297 ml/min clean air to the air stream, thus diluting by another factor of 100. (*b*) *Top*: Raw PID signal to 5 consecutive fluctuating 2-heptanone stimuli, produced using a mineral oil dilution (*left*) and an air dilution (*right*). The strength of the PID signal is proportional to odorant concentration. *Bottom*: Mean PID signal during 0.5-2.0 s, divided by the first trials’ value. In the mineral oil dilution system, the odorant concentration decreases from trial to trial, but in the air dilution system, the odorant concentration remains stable.

**Figure S2.**
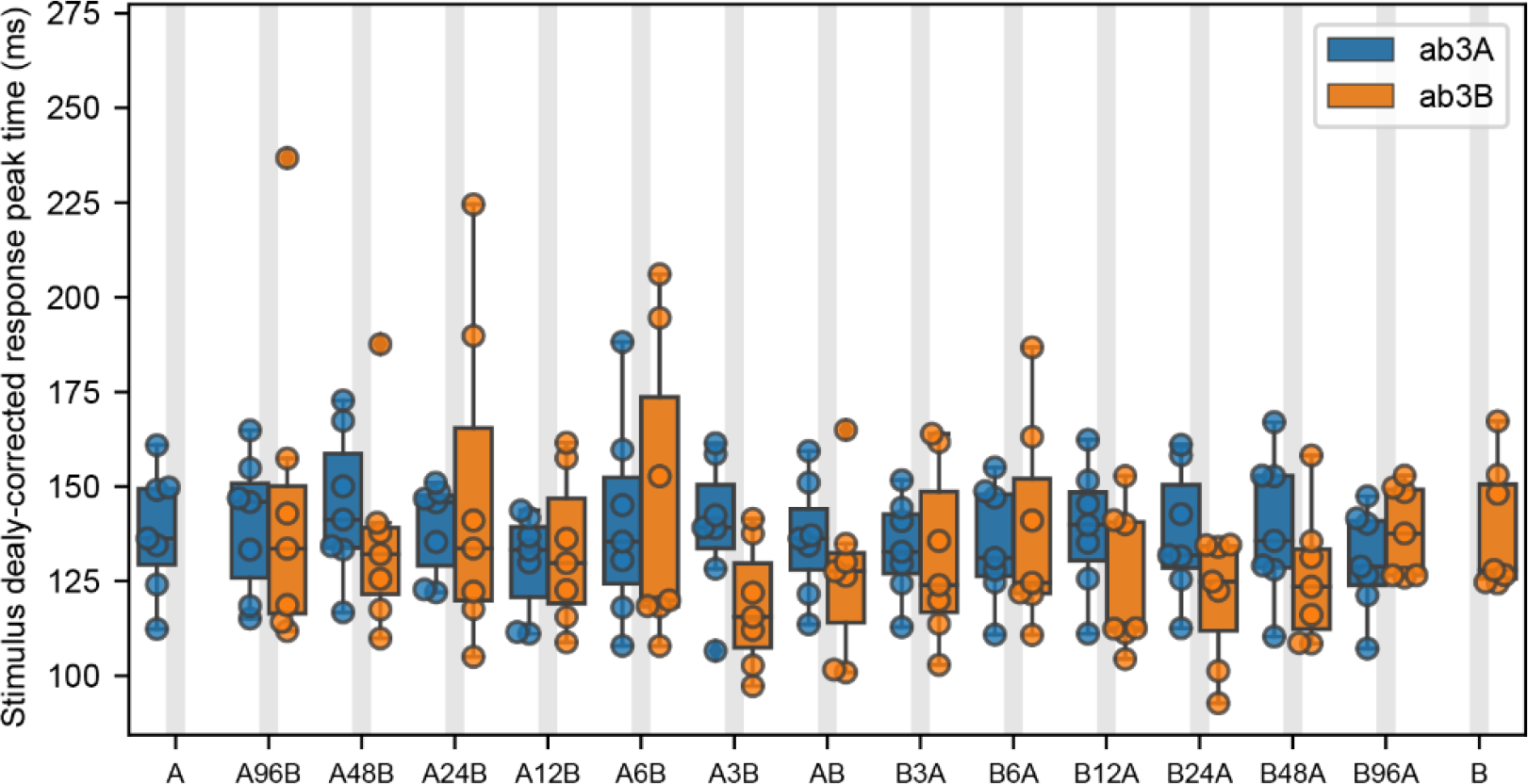
Ephaptic inhibition does not affect the timing of odor responses. Same data as in Fig. 3c, but the response peak timing was analysed. N = 7 sensilla in 7 flies. Horizontal lines show medians, boxes show interquartile ranges, and whiskers extend to ±1.5 times the interquartile ranges.

